# Paired Associative Stimulation Fails to Induce Plasticity in Freely Behaving Intact Rats

**DOI:** 10.1101/765636

**Authors:** Windsor Kwan-Chun Ting, Maxime Huot-Lavoie, Christian Ethier

**Author notes:** **Corresponding Author: Dr. Christian Ethier, Centre de Recherche CERVO, Département de psychiatrie et de neurosciences, Université Laval**, 2601 Chemin de la Canardière, Québec, QC; G1J 2G3.

## Abstract

Paired Associative Stimulation (PAS) has been explored in humans as a non-invasive tool to drive plasticity and promote recovery after neurological insult. A more thorough understanding of the phenomenon of PAS-induced plasticity is needed to fully harness it as a clinical tool. Here, we tested a PAS model in awake rats in order to study the principles of associative plasticity. Using chronically implanted electrodes in motor cortex and forelimb, we explored PAS parameters to effectively change corticomotor excitability, assessed using closed loop, EMG-controlled cortical stimulation. We tested eleven stimulation intervals, chosen to force the coincidence of neuronal activity in the rats’ motor cortex and spinal cord, with timings relevant to the principles of Hebbian spike-timing-dependent plasticity. However, despite a relatively large number of stimulus pairings (300), none of the tested intervals reliably changed corticospinal excitability relative to control conditions. Our results question PAS effectiveness under these conditions.

## Introduction

### Spike Timing as a Driver of Synaptic Plasticity

Donald Hebb first postulated that coincident activity between active neurons could induce a change in the strength of their connecting synapses (Hebb 1949). The propensity to strengthen or weaken synapses based on coincident activity could be a model of learning, paving a mechanistic foundation for connecting biology with behaviour. At the outset however, it was clear that Hebbian learning principles would need to be further refined to accurately describe biological phenomena at the synapse. Seminal studies in spike-timing-dependent plasticity (STDP) (Markram, Lubke et al. 1997, Bi and Poo 1998) have been conducted to develop models of this plasticity at the synaptic level. Whether synaptic potentiation (Long Term Potentiation, LTP) or depression (Long Term Depression, LTD) occurs is contingent upon the pattern of firing activity in the pre- and post-synaptic neurons (Cooper 2005).

The Hebbian STDP model posits that activity in the pre-synaptic neuron should occur prior to activity in the post-synaptic neuron for facilitation to occur at the target synapse (Feldman 2012). One advantage of the Hebbian STDP model is that accidental coincidences in pre- and post-synaptic firing do not significantly change the strength of the synapse, and uncorrelated activity will lead to a weakening of the connection (Song, Miller et al. 2000). It suggests a method of neuromodulation, making conceivable the artificial restructuring of cortico-cortical and cortico-spinal circuits involved in motor control towards more beneficial and functional connections by inducing repetitive paired activity at a desired set of synapses. This may even represent a framework for harnessing beneficial plasticity after cerebrovascular or traumatic insult.

However, the STDP hypothesis as it applies to larger circuits such as the corticomotor system is contingent upon certain assumptions, one being that principles derived from *in vitro* studies at the synaptic level remain sound when applied to higher level systems *in vivo*. Beyond the complexity of the system’s anatomy, ongoing patterns of neural activity (spontaneous or behaviour related) may interfere with the fine-tuned firing patterns which STDP putatively requires. Hence, although STDP has been well-studied at the synaptic scale, to really harness this model effectively, we need to study it more systematically at the systems level in conjunction with ongoing neuronal activity.

### Non-invasive Paired Associative Stimulation (PAS) in Humans

Paired Associative Stimulation (PAS) is a non-invasive method of modulating the excitability of the motor cortex and downstream corticospinal connections to facilitate the recruitment of targeted muscles, based on Hebbian STDP principles. The idea is that repeated coincidences of stimulation-induced activity in targeted areas promotes strengthening of useful neural connections within the sensorimotor system.

The first clear demonstration of PAS in humans was designed to promote plasticity at the cortical level (Stefan, Kunesch et al. 2000). Stefan et al. timed Transcranial Magnetic Stimulation (TMS) with the cortical arrival of afferent activity induced by electrical stimulation of the median nerve, which elicited an increase in cortical excitability. This was assessed by comparing resting motor evoked potential (MEP) amplitude from TMS stimulation in the abductor pollicis brevis (APB) muscle of healthy humans, before and after intervention. The results indicated that topographically specific and sustained (30-60 min) increases in excitability of the motor system was possible through non-invasive PAS in man, by carefully timing cortical stimulation with somatosensory signals afferently propagated towards the cortex.

Taylor and Martin performed the first study demonstrating in humans that PAS could also be used to induce a plasticity in the spinal circuits (Taylor and Martin 2009). They demonstrated that by timing peripheral nerve stimulation so that antidromic potentials in motoneurons reached the cell bodies in the spinal cord one millisecond after the arrival of TMS-induced corticospinal volleys, they could induce a sustained potentiation of the corticomotor circuits controlling the biceps brachii muscle. Since that time, several studies have attempted to validate this phenomenon with mixed success, and studied human PAS to target plasticity in the neural circuits controlling upper and lower limbs of man; see (Carson and Kennedy 2013, Suppa, Quartarone et al. 2017) for reviews on the subject.

PAS is seen as a potential tool to strengthen residual circuits after neurological injury and promote functional recovery. In humans, PAS has been studied as a potential therapeutic intervention. Bunday et al. (2012) studied the potential to use PAS to reshape functional corticospinal connections in human patients with spinal cord injury (Bunday and Perez 2012). Pairing TMS over the motor cortex and Peripheral Nerve Stimulation (PNS) of the ulnar nerve, they were able to potentiate the MEPs and maximal voluntary contraction force obtained in the first dorsal interosseous muscle using an STDP-based protocol. The effectiveness of a subsequently modified PAS protocol in increasing voluntary force output was replicated in the tibialis anterior hindlimb muscle of the participants as well (Urbin, Ozdemir et al. 2017). Subsequent research has extended modified PAS protocols with mixed success in improving functional recovery after neurovascular insult, both in animals (Shin, Han et al. 2008) and in humans (Castel-Lacanal, Gerdelat-Mas et al. 2007, Castel-Lacanal, Marque et al. 2009, Rogers, Brown et al. 2011, Cho, Sabathiel et al. 2016, Ferris, Neva et al. 2018, Palmer, Wolf et al. 2018, Tarri, Brihmat et al. 2018). PAS initially showed great promise for rehabilitation, however enthusiasm for this approach has been dampened by inconsistent results. PAS was shown to have a very high inter-subject variability (Sale, Ridding et al. 2007, McGie, Masani et al. 2014, Tarri, Brimhat et al. 2018), its effects are strongly dependent on mindful, persistent attention on the target limb (Stefan, Wycislo et al. 2004) or even failed to induce any consistent plastic effects (McGie, Masani et al. 2014).

#### Invasive PAS in Animals

To better characterize the effects of PAS and clarify its potential as a therapeutic tool, research groups have more recently begun investigating this approach in animal models. Mishra et al. (2017) performed cervical epidural stimulation in conjunction with stimulation in the rodent motor cortex, to create sustained increases in biceps MEPs arising from cortical stimulation (Mishra, Pal et al. 2017). However, these experiments were done under anaesthesia, leaving open the possibility of pharmacological effects of Ketamine/Xylazine preparations (Yang, Chang et al. 2011, Huang and Yang 2015) and thereby reducing the ecological validity of the results. Non-invasive PAS in restrained, awake rats can drive potentiation of the *hindlimb* corticospinal circuits using stimulation intervals between cortex and tibial nerve of 5 and 15 ms (Zhang, Sui et al. 2018). A non-invasive model under anaesthesia has also been used to investigate PAS for therapeutic improvements after stroke, using TMS over the rat cortex and coincident peripheral nerve stimulation using electrodes inserted into the contralateral soleus hindlimb muscle (Shin, Han et al. 2008), finding that specific interstimulus-intervals could potentiate MEP amplitudes. Clearly, there has been a convergence of research in recent decades, from both clinical and fundamental sciences towards applying timed stimulation to the motor system to promote plasticity.

### Objective

The project aimed to establish an effective animal PAS paradigm in the *forelimb* using an awake and invasive rat model, to better understand the fundamental mechanisms of associative plasticity and ultimately to improve interventions in humans for neurorehabilitation.

### Hypotheses and Predictions

We hypothesized that we could use PAS to modulate corticomotor excitability toward target contralateral muscles in the forelimbs of rats. Further, based on Hebbian principles we predicted that LTP-like plasticity would be induced with spike timing conditions where the pre-synaptic descending volley arrived at the level of the spinal cord a few milliseconds *prior to*, or *at the same time as* the antidromic post-synaptic volley arising in motoneurons from peripheral stimulation of the target muscles. By holding other PAS parameters constant (stimulation amplitude, frequency, number of pulses), we tested PAS protocols in a randomized trial and predicted that different stimulation intervals would induce systematic changes in corticomotor excitability as predicted by the STDP model.

## Results

### Failure of Spike Timing to Modulate Cortical and Spinal Plasticity

We tested a wide range of STDP-relevant intervals between cortical and peripheral stimuli (ISI conditions), in a randomized fashion for each rat and used an EMG-controlled closed-loop method to measure pre- and post-intervention MEPs. We found no significant modulation of corticospinal excitability using our PAS intervention *in vivo*. We analyzed the MEP amplitudes obtained from cortical stimulation probes before and after each PAS intervention using a mixed model ANOVA on the normalized data. Probe time (SESSION) was considered a repeated-measures fixed factor, and ISI Condition (CONDITION) was the second fixed factor. We included a SESSION x CONDITION interaction term in the statistical model, and a random factor for the rat accounting for the randomized design. We did not find a significant effect of our PAS intervention for any of the timings we tested (Figure 2). Statistically, there was no significant interaction between SESSION (pre- and post-intervention MEPs) and CONDITION (F (28, 245) = 0.53, p > 0.05), Table 1), meaning that there was no ISI condition for which the intervention resulted in a statistically significant change in corticospinal excitability over assessment time. There was a significant main effect of ISI Condition however, F (14, 248) = 1.74, p = 0.049), independent of the time at which the MEP was measured post-intervention, indicating that the ISI condition had a barely significant effect on the MEP amplitude. However, a two-way Dunnett’s test used *a posteriori* to make the relevant pairwise comparisons between the no stimulation control condition and all other STDP and PAS control conditions, while controlling for family-wise error rate at 5% did not detect significant differences. There was no significant main effect of SESSION, F (2, 245) = 0.12, p > 0.05. In summary, our statistical analyses did not support the efficacy of PAS under these conditions.

**Table 1:** Mixed Effects Model Analysis, Fixed Effects (Type III) Summary using the Restricted Maximum Likelihood Estimation (REML) Method.

| Term | DF Num | DF Den | F-Value | P-Value |
| --- | --- | --- | --- | --- |
| Session | 2.00 | 245.42 | 0.12 | 0.886 |
| Condition | 14.00 | 247.91 | 1.74 | 0.049 |
| Session*Condition | 28.00 | 245.42 | 0.53 | 0.975 |

### Control Experiments

Four different control protocols where we did cortical/muscle stimulation in isolation, no stimulation, and maintained a large offset between paired stimuli, respectively did not significantly alter corticomotor excitability (Figure 2, conditions to the right of vertical dotted line). Interestingly, we noted a trend towards a depressive effect for the cortical stimulation only (mean ratio post/pre = 0.86), muscle stimulation only (0.93), and ISI +505ms stimulation (0.85) conditions, but not in the control condition for which rats received no stimulations at all in place of a PAS intervention (0.98).

## Discussion

There exists a mixed literature on human PAS and several variations of the original protocol (Stefan, Kunesch et al. 2000), with some convincing reports demonstrating its effectiveness at inducing at least transient changes in corticomotor excitability (Taylor and Martin 2009, Bunday, Urbin et al. 2018), and others showing ineffective interventions or highly subject-dependent results (Muller-Dahlhaus, Orekhov et al. 2008, McGie, Masani et al. 2014). Our own results support the latter findings. Our PAS protocol, with parameters inspired by typical interventions in humans, was ineffective at modulating plastic changes in corticospinal excitability. There was no significant interaction between fixed factors, and the critical comparisons between the hypothesized potentiating ISI conditions with the no stimulation control condition were all non-significant, leading us to conclude that our PAS protocol was ineffective overall in potentiating corticospinal connections.

### PAS Parameter Space

Setting aside spike timing, the entire parameter space for a PAS intervention protocol is vast, with no known physiological principles guiding a specific combination of stimulation intensity, frequency and/or number of repetitions over another. Consistent with most studies, we chose above-threshold but sub-maximal stimulation amplitudes (1.5x and 1.25x motor threshold for muscles and cortex respectively). We decided on a PAS protocol with a number of paired stimulations (300) and stimulation frequency (0.5 Hz) on the higher end compared to most other published protocols (Suppa, Quartarone et al. 2017). We reasoned that if anything, this would enhance any PAS effects. It would be possible but counterintuitive that these differences reduced the likelihood of inducing plastic changes.

### Neuromuscular Fatigue

We observed that MEPs after PAS interventions were generally smaller than the average of the baseline measurements. This trend was surprising given the evidence that single pulses of peripheral electrical stimulation are insufficient to change corticomotor excitability with or without coincident voluntary contraction in humans (Saito, Sugawara et al. 2014), and that higher frequencies are needed for supraspinal effects (Grospretre, Gueugneau et al. 2017). This trend was also present for control conditions involving cortical or muscle stimulation alone, but not when the rats received no stimulation in place of PAS. These results indicate that the probes themselves had no effects, but that all stimulation interventions induced a low level of spinal and/or supraspinal fatigue, in a manner similar to that of a sustained fatiguing muscle contraction, which has been shown to evoke a reduction in TMS-induced MEP size (McKay, Tuel et al. 1995, Pitcher and Miles 2002). Finally, the physiological recruitment order of motoneurons using electrical stimulation is also inversed compared to natural activation patterns (Hennings, Kamavuako et al. 2007), so this may have contributed to the fatigue-like responses we observed throughout.

### Closed-loop Assessment

Another plausible explanation for our negative results is the high intrinsic variability observed in the MEP responses of our rats during free behaviour. While we attempted to reduce this through our closed-loop stimulation design, enabling us to probe the corticomotor system only when the pooled motoneuron output (EMG) was within a fixed range of activity, it nevertheless remained high. Our EMG-based closed-loop pre- and post-intervention assessment probes were specifically designed to assess the excitability of the corticomotor system at relatively similar, low levels of EMG activity (approximately 5-15% of maximum EMG amplitude observed under free behavior). The aim was to minimize MEP variability, by avoiding stimulating in different conditions of corticomotor excitability, such as during a strong voluntary contraction or during reciprocal inhibition acting on the recorded muscle. However, the PAS intervention itself was not completed in an EMG dependent manner, because we could not record and stimulate muscles simultaneously with our setup. Perhaps applying the closed-loop approach to the paired stimulation as well, would have allowed for a more systematic and reproducible recruitment of neuronal elements, thereby leading to more reliable PAS effects.

### Stimulation Models and Specificity

In our chronic PAS model, we inserted electrodes directly into the target muscle, and validated this approach in an acute experiment to verify that electrical stimulation of the muscle fiber was sufficient to generate antidromic volleys back-propagating to the deafferented spinal cord. Compared to direct nerve stimulation, intramuscular stimulation may result in a small difference in the relative timing of stimulation-induced antidromic motoneuron activation and orthodromic afferent activity. In addition, direct nerve stimulation can recruit a larger number of fibers of all modalities, not limited to a specific target muscle, but including all motor and sensory fibers traveling in the nerve at the chosen stimulus location. These differences in peripheral fiber recruitment may have contributed to the apparent inconsistency between our results and that of others showing the effectiveness of PAS using electrical stimulation in rodents (Mishra, Pal et al. 2017).

With respect to the PAS literature, we can hypothesize there may be intrinsic differences in MEP variability between non-invasive (TMS, the standard technique for PAS in humans) and invasive (Intracortical Microstimulation, ICMS) stimulation methods due to different circuits being recruited. ICMS, although having a greater spatial and temporal resolution than non-invasive methods of neural activation such as TMS, is also non-specific in the sense that it activates all types of neurons and other cell types. The exact recruitment patterns of the cortical circuits are of very little theoretical importance for PAS interventions targeting spinal circuits, as long as a corticospinal volley occurs in a timely manner with that of peripheral stimulation. Therefore, especially since it was previously used successfully in rats (Mishra, Pal et al. 2017), it would be surprising if our use of ICMS was factor explaining our negative results.

Some electrophysiology studies have suggested there are both monosynaptic and polysynaptic connections onto rats’ corticospinal motoneurons (Elger, Speckmann et al. 1977, Liang, Moret et al. 1991, Hori, Carp et al. 2002), but more recent work has suggested the rat corticospinal tract is exclusively polysynaptic (Alstermark, Ogawa et al. 2004). Although this is a major physiological difference between rats and primates, we believe that this is not a critical factor to explain differences between our negative results and successful human PAS. In addition to the variable conduction time between individual fibers, a polysynaptic pathway will increase the temporal spread of action potentials in a stimulation-induced volley. The ascending afferent sensory pathway in humans is polysynaptic (Abraira and Ginty 2013), and yet PAS is still effective when TMS is timed with the arrival of afferent volleys in the cortex (Stefan, Kunesch et al. 2000). By the same token, we expected that a polysynaptic descending pathway would not prevent us from timing the descending volley with antidromic motoneuron activation.

### Opposing Plastic Changes Along the Corticomotor Pathway

Thinking along these lines, however, the ISI timing offset of the paired stimulation dictates the target location of plasticity. In an ideal world, the effects will be localized only to one target area. However, since the corticomotor contains multiple synaptic connections, any given ISI condition predicted to induce LTP-like changes at one site according to Hebbian STDP (the motor cortex for example), could lead to LTD-like effects at the second site (the spinal cord for instance), and vice versa. In rats, we estimated the interval between PAS-induced pre- and post-synaptic activity at the cortex and the spinal cord for given cortical and peripheral stimulation intervals. These opposing effects are reflected in the lack of situations where LTP-like effects can be predicted at both spinal and cortical levels, represented by numbers in green (Figure 2). This competition between potentiation and depression at different locations may reduce the PAS effectiveness to induce a net increase in corticospinal excitability. Due to the non-invasive nature of human PAS experiments, this can potentially be an explanatory factor for the variance in PAS effectiveness observed in the clinical data. This issue can be dissected in animal models with terminal *ex vivo* experiments, but addressing this issue *in vivo* will require advances in our stimulation methods to be simultaneously non-invasive, yet highly spatially specific. The goal here would be to isolate the bookends of the paired stimulation just bounding the targeted synapses. That would be a seminal advance in addressing the utility of PAS *in vivo*.

### Seeking the Perfect Storm

Voluntary effort itself has been shown to be a necessary driver for potentiation in humans for specific PAS protocols (Kujirai, Kujirai et al. 2006), with two proposed mechanisms being the reduction of intracortical inhibition networks coincident with contraction, or the facilitatory effect of attention via the activation of memory systems (Stefan, Wycislo et al. 2004), but this is contradictory to earlier cited findings that PAS works well under anaesthesia in animals (despite the differences among species). The effect of known neuromodulators on PAS, such as dopamine, should not be underestimated particularly because of its direct role in mediating neuronal potentiation (Yagishita, Hayashi-Takagi et al. 2014), and its broader implications in maintaining attention (Suppa, Quartarone et al. 2017). Additional factors influencing PAS effectiveness are numerous, and may include even the time of day – a study in humans showed that PAS sessions performed in the afternoon were significantly potentiated in one study, whereas sessions completed in the morning did not (Sale, Ridding et al. 2007). In that paper, variance was attributed to circadian effects and specifically the inhibitory effect of cortisol on plasticity. These examples drive home the point that our knowledge of what coincident factors are required to induce LTP-like potentiation remains limited, and based on our study, future studies should likely *not* be restricted to simple application of Hebbian principles; it may not be enough.

PAS has also been reported to exhibit high variance depending on the subject being tested. McGie et al. (2014) conducted a non-invasive PAS study in humans, with the goal of comparing different paired stimulation protocol frequencies (McGie, Masani et al. 2014). Tarri et al. (2018) studied the effect of PAS in humans as a therapeutic adjunct to stroke using a randomized double-blind controlled approach (the CIPASS Trial) (Tarri, Brimhat et al. 2018). Both groups reported high between-subject variability in PAS outcomes but found no consistent effect of PAS targeting spinal circuits, attributing the variability observed to individual factors such as the lesion size / location, and different rehabilitation intensiveness, both influencing the physiological capacity available for PAS effects. Importantly, the degree of muscle facilitation can vary greatly even *within the same participants* across repeated PAS sessions (Tarri, Brimhat et al. 2018). These studies emphasize the mercurial nature of PAS effectiveness even within individuals, and the highly stereotyped/specialized conditions necessary for consistent beneficial effects to become apparent. It may turn out that a conjunction of multiple concurrently acting factors is necessary in order to facilitate PAS potentiation under free behavior in animals.

### Conclusion

In conclusion, our data does not support the effectiveness of PAS in promoting plasticity through the Hebbian STDP model in freely behaving rodents. Our initial goal was to develop a clinically relevant animal model for paired stimulation which would have allowed more detailed studies and optimize interventions. Although the model itself was developed successfully, this series of experiments suggested that an open-loop PAS intervention in a freely moving animal is not effective to reliably drive plasticity in the corticospinal system. Our results highlight the complexity of associative plasticity and demonstrate that forced coincidence of neuronal activity is not sufficient to reliably potentiate corticospinal excitability. Future research will need to investigate whether other variations in the PAS parameter space, reduction of interference from ongoing neuronal activity or manipulations of neuromodulators may be required to drive corticospinal potentiation more reliably. This will determine whether PAS indeed has potential as an interventional measure for modulating corticomotor plasticity.

## Methods

### Animals and Surgical Preparation

All animal handling and experimentation were performed in accordance with institutional review board (Comité de Protection des Animaux de l’Université Laval), and guidelines from the Canadian Council on Animal Care. Nine Long Evans rats and one Sprague-Dawley rat (all male) were housed under 12/12 inverse light-day cycle with food and water available *ad libitum*. 150 experimental sessions were planned in ten animals, to investigate the effectiveness of 15 inter-stimulus intervals (ISI), including controls. However, due to rare implant failure, some ISIs were not tested in all the rats. All ISI conditions were tested in a minimum of five animals, and an average of seven to eight (Supplementary Table 1). Our PAS intervention typically targeted the *extensor carpi radialis* muscle. However, in some animals, we used pairs of EMG wires implanted in more proximal locations (*Biceps* or *Trapezius muscles*). The distribution of muscles tested within the full dataset is shown in Supplementary Table 2.

### Chronic PAS Implantation Surgery

During aseptic surgeries performed under isoflurane anesthesia, rats were implanted with 1 mm x 1 mm custom square arrays of four 80/20 platinum-iridium electrodes, each 75 microns in diameter. The array was inserted into the caudal forelimb area (CFA) of the primary motor cortex (M1) by a stereotaxic craniectomy, centered at 1mm anterior and 3.5mm lateral relative to bregma, and the surrounding exposed dura matter was covered with silicone gel for protection. Three pairs of PFA-coated multi-stranded stainless-steel wire electrodes (A-M Systems Inc.) were inserted into the contralateral *extensor carpi radialis (ECR)*, *biceps brachii* and *trapezius* muscles (the latter two serving as alternative muscles in case the ECR electrode failed). EMG and cortical electrodes were pre-soldered to either an InVivo1 MS12P or a SAMTEC 2×7 connector, which were secured to the skull with dental cement and six bone screws as anchors. A posterior skull screw served as the ground electrode for cortical monopolar stimulation. An additional reference electrode for the EMG measurement was embedded subcutaneously in the upper back. See the Figure 1 inset for a visual schematic of the electrode array as well as the implant location. Animals recovered undisturbed for a week after implantation prior to testing and were given time to familiarize themselves with being connected prior to data collection.

**Figure 1:**
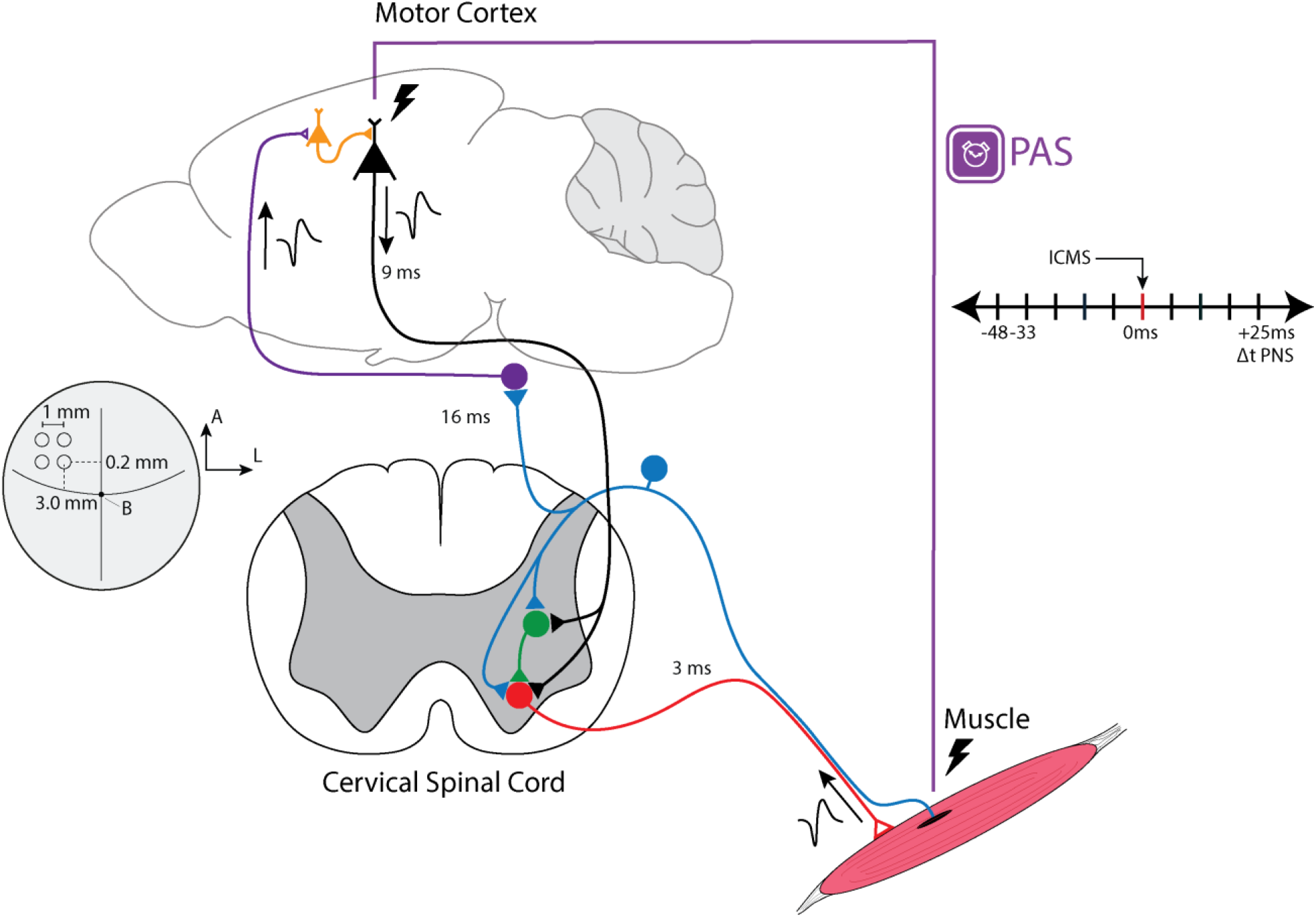
Spinal Cord and Cortical PAS. To induce plasticity at the level of the spinal cord, we timed the descending action potentials from cortical ICMS to arrive at the spinal cord with a set latency before or after the arrival of the ascending antidromic action potentials coming from peripheral stimulation. We know from our and previous studies under anaesthesia that it takes approximately 9 ms and 3 ms for both signals to arrive at the spinal cord, respectively. To induce plasticity at the level of the cortex, we stimulated the periphery first and waited for 16 ms for the signal to cause pooled action potentials in the motor cortex. Timing offsets for subsequent cortical stimulation were calculated based on these conduction latencies. **Inset:** Dorsal view of the rat brain, showing where we inserted the platinum-iridium electrode array in the caudal forelimb area (CFA) of M1. The medial posterior electrode was positioned 3.0 mm ML (left), 0.2 mm AP, and −1.5 mm DV. All other leads were 1 mm anterior (A) and lateral (L) from this reference electrode, as illustrated.

**Figure 2:**
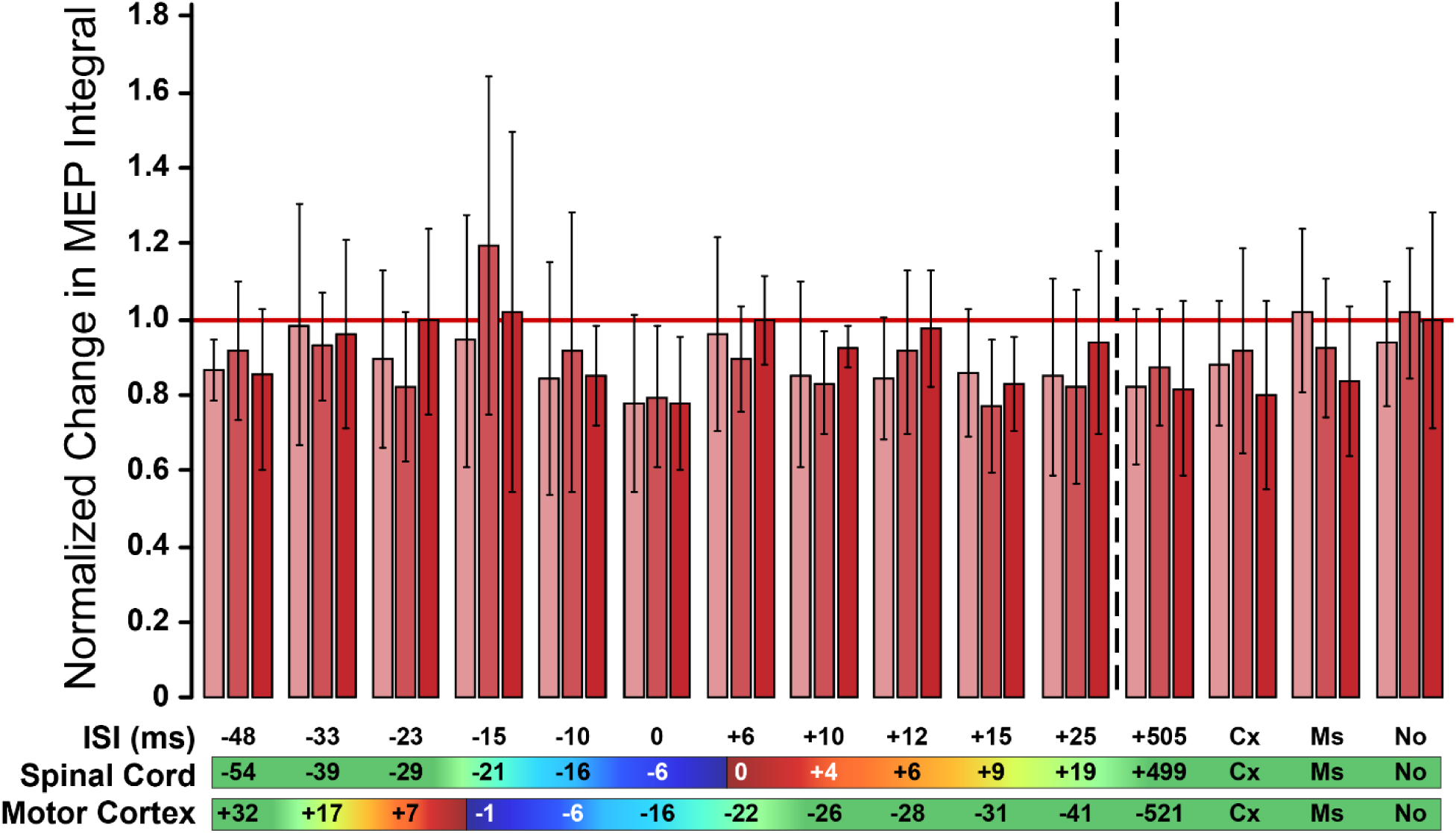
PAS does not significantly potentiate MEP responses *in vivo.* Main effect plot, depicting that for Post 1 (2 min after PAS), Post 2 (17 min after PAS), and Post 3 (32 min after PAS) sessions, there were no significant differences between STDP experimental conditions and control conditions. Error bars are 95% Confidence Intervals about the mean. The horizontal reference line marked in red signifies no change between that post condition and the baseline average. Control conditions are shown to the right of the vertical dotted line, to separate them from the ISI conditions tested to the left (Cx = Cortical Stimulation Only, Ms = Muscle Stimulation Only, No = No Stimulation). Stimulation (ISI) Offset refers to the latency between stimulation of the Cortex and the Muscle. Positive numbers refer to muscle stimulation occurring after cortical stimulation. Delta Spinal Cord is the estimated latency between the arrival of the descending volley onto the motoneurons in the spinal cord and the arrival of the antidromic action potentials evoked from muscle stimulation; Delta Motor Cortex is the estimated latency between the arrival in M1 of peripheral stimulation-induced afferent activity and the motor cortex stimulation. Red indicates conditions we predicted to induce LTD-like effects, green indicates PAS conditions we predicted would induce LTP-like effects.

### Study Design and Experimental Paradigm

We used a repeated-measures randomized block design (same rat tested on all ISI conditions in a randomized order) to test the effect of STDP Timing Condition on the change in integral of the averaged MEP response after the PAS experiment.

We tested ten rats with chronic implants. Each rat was to be tested once in each condition. To account for possible order effects inherent to a within-subjects design, the order of testing conditions was randomly assigned using the *randperm* function in MATLAB. One condition (ISI −15ms) was added at the end for six rats which had all data collection completed, based on another study (Zhang, Sui et al. 2018) which showed a promising timing condition and was published while data collection was in progress. For rats with whom data collection had not started yet, a re-randomization was performed to integrate this new condition. For each rat, each test was separated by roughly 24 hours, to minimize carryover effects between previous paired stimulation interventions.

We tested four control conditions: three PAS controls, involving (1) cortical stimulation only, (2)-peripheral stimulation only, and (3) no stimulation, as well as (4) one extra-long ISI timing control involving paired stimulation of the motor cortex and the contralateral peripheral muscle offset by +505 ms. We reasoned that if timing during paired stimulation was the driving factor behind plasticity, and not the pairing of stimulation *per se*, this condition should have a null effect comparable to the previous PAS control conditions.

Each experiment followed a fixed schedule (see Figure 3). After connecting the rat to the hardware interface, we completed three 5 minute “probes” to assess the corticomotor excitability prior to the PAS intervention (see following section). The probe was completed when 30 stimulations were delivered, or 5 minutes had elapsed, whichever came first. Probes were separated by 10 minutes each. After each PAS intervention, three post-PAS probes were completed in the same manner to assess excitability of the corticomotor system after paired stimulation. We allotted 2 minutes for wire switching and software changes, immediately before and after each PAS intervention.

**Figure 3:**
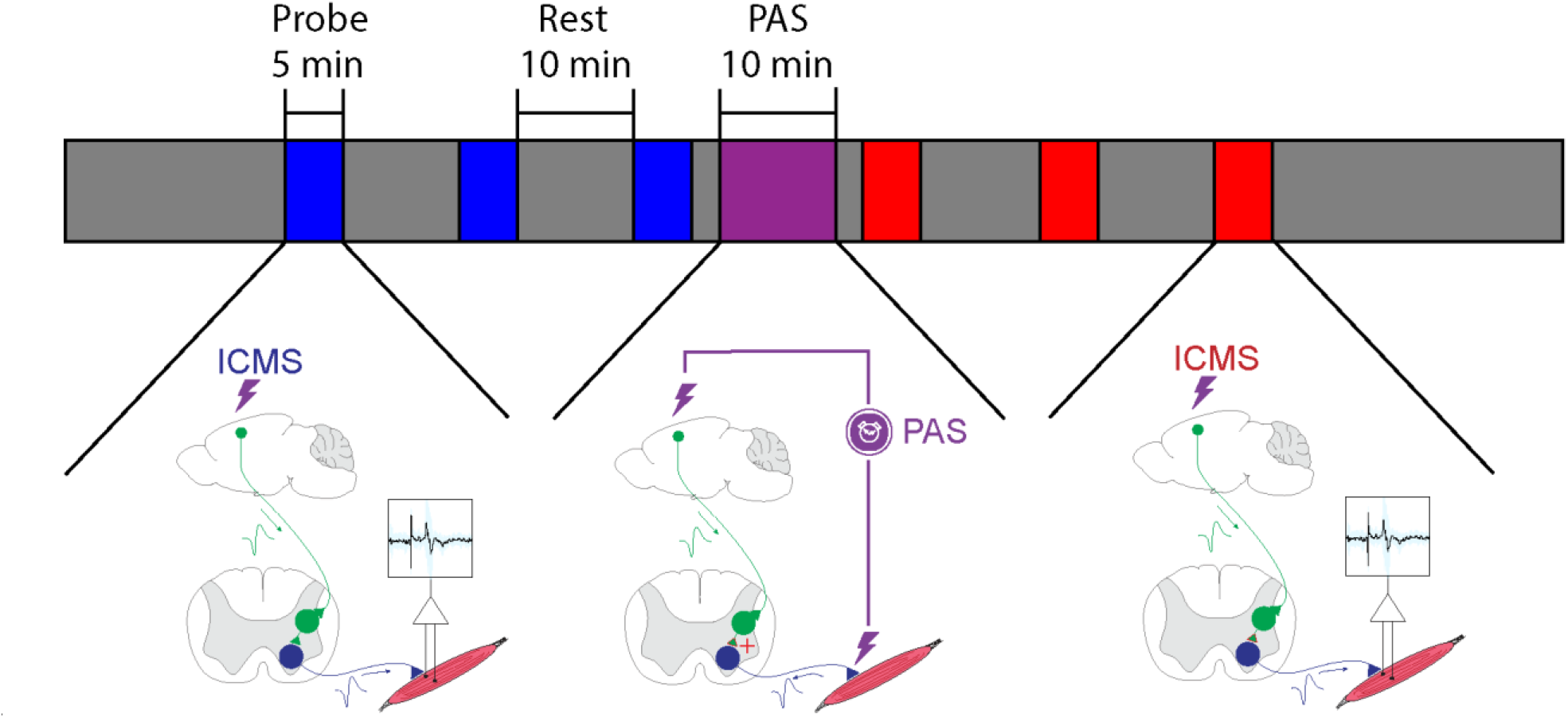
Experimental Design. A: Each session began with three 5-minute probes in which we performed closed-loop EMG-dependent cortical stimulation to assess baseline MEPs amplitude, each separated by 10 minutes. The PAS session itself, involving 300 pairs of stimuli to the cortex and the muscle at a rate of 0.5 Hz, took about 10 minutes. This was followed by three post-PAS probes so we could assess corticospinal excitability up to 30 minutes after paired stimulation for each inter-stimulus interval. After each experiment, we manually verified all MEPs using custom software and excluded traces with movement artifacts or noisy EMG signals.

### Probe Assessment of Corticomotor Excitability

To assess corticomotor excitability before and after PAS, we compared the size of MEPs obtained from cortical stimulation using a closed-loop stimulation protocol. The EMG activity in the target muscle was continuously measured, and cortical stimulation was triggered in real-time if the activity reached within [2 12] SD above the baseline (defined as the mean value of the rectified EMG signal measured over two seconds when the limb was fully relaxed), and the if EMG activity was on the rising phase (contraction was being initiated during free behaviour as opposed to when the muscle was relaxing from a previously larger contraction). We thought that restricting the conditions for stimulation to a low-moderate level of voluntary activation of the corticospinal system achieved during free behavior (walking/grooming/exploring) would reduce the baseline MEP variability observed with cortical stimulation (Darling, Wolf et al. 2006). To do this, we used the *envelope* function in MATLAB, which calculated the peak envelope of the filtered data with a moving spline over the local maxima of the previous 32 data points. Cortical stimulation was contingent upon the EMG envelope crossing the pre-determined activity threshold. The variability was still high albeit reduced after applying the closed-loop stimulation protocol, so we averaged all three baseline measurements during the statistical analysis to obtain an overall assessment of corticomotor excitability prior to the PAS intervention. We recorded from different muscles, depending on the location where we obtained the best quality MEPs (Supplementary Table 2). We always started with the most distal forelimb muscles (the extensors) and moved proximally for each rat, as necessary. In one rat, we used a monopolar EMG recording configuration resulting in an EMG signal contaminated with crosstalk from cardiac activity. We manually adjusted the upper and lower limit of the EMG window enabling probe stimulation, in a manner that better reflected a low amplitude muscle contraction.

### Electrophysiological Data Acquisition and Stimulation Configuration

Independent paired electrical stimulation protocols were achieved through two A-M Systems (Sequim, WA, USA) 2100 stimulators, each connected to separate pins on an *InVivo1* (Roanoke, VA, USA) commutator through a custom-made breakout board interface. Multi-channel recording was made possible by routing the EMG signal into a Brownlee Precision Model 440 (Santa Clara, CA, USA) Instrumentation Amplifier. Within this unit, a signal gain of 100, a bandpass filter between 50 Hz and 1.0 kHz was used for EMG, and bandpass of 1-300Hz for LFP signals. A 60Hz notch filter was applied. This output signal was split in two, with one copy being routed into a Powerlab 8/sp unit by AD Instruments (Colorado Springs, CO, USA), and further processed with a 10 Hz highpass filter before being saved. The second copy was routed into a National Instruments (Austin, TX, USA) Digital to Analog Converter (DAC) SCB-68A system, which was operated via custom MATLAB software. We used the DAC system and MATLAB software to initiate all probe and PAS stimulation protocols, via the trigger input ports on the A-M systems stimulators.

### Latency Measurements and Sign Convention for Spike Timing Experiments

To confirm the conduction latencies, we completed a series of acute experiments under ketamine/xylazine and urethane anaesthesia, in rats with a similar weight and size to those used for chronic implants. We performed an acute laminectomy under urethane anaesthesia to measure the antidromic conduction time in motoneurons between the muscle and the spinal cord. We exposed the dorsal spinal cord between the C4 and C6 regions. With the dura intact, we inserted a tungsten electrode, 127 um in diameter in the C5 region ipsilateral to the right forelimb, 1.0 mm lateral to the midline. We also inserted a pair of EMG electrodes in the right *extensor carpi radialis* using the same method as in the chronic implants. Stimulation of the spinal cord C5 region using single pulses led to an isolated wrist extension in the rat’s forelimb, verifying the location of the ECR motoneuron pool for efferent connections (Tosolini and Morris 2012). The dorsal spinal cord was deafferented between C3 and C7, to isolate antidromic propagation instead of conduction along afferent sensory fibers. Following this, we stimulated the EMG electrodes and recorded local field potentials (LFPs) from the electrode site in C5. Filtering parameters for the LFP recording included a bandpass of 1-300Hz, with a 60Hz notch filter applied and a gain of 100. Data was averaged across 200 stimulations. We determined the afferent latency to be 3 ms (Supplementary Figure 1A).

To measure the time a neuronal volley requires to reach the cortex after muscle stimulation, we recorded local field potential (LFP) responses in M1 following intramuscular stimulation (Supplementary Figure 1B). We postulated that the peak of the initial negative inflection in the local field potential from contralateral muscle stimulation reflected the time at which the greatest neural activity is observed amongst the post-synaptic neurons in the cortex. We inserted a pair of EMG electrodes in the right *extensor carpi radialis*, then inserted one platinum-iridium electrode 1.5 mm DV into M1 centered at the array coordinates for the other rats. A second electrode about 1 mm lateral from the first was positioned on the surface of the dura. Both cortical electrodes were connected by a common ground at the skull screw, and local field potentials (LFPs) were measured by calculating the voltage differential between the cortical electrodes with the same LFP recording parameters above. Stimulation was delivered to the EMG electrode in the right ECR through bipolar single pulses with a 0.2 ms duration, repeated at 0.5 Hz. Data was averaged across 360 stimulations. The afferent latency from ECR stimulation to cortical evoked potential was 16 ms (Supplementary Figure 1B). With an average MEP latency of 12 ms for ECR and a 3 ms travel time from muscle to spinal cord, we estimated a latency of 9 ms for a cortical stimulation-induced descending volley to reach motoneurons, including synaptic integration time. These latencies are reported in Figure 1.

Using these conduction latencies, we chose a set of stimulus intervals which would result in various pre- and post-synaptic timing relevant to the rules of spike-timing-dependent plasticity, either at the cortical and/or spinal levels. The full list of ISI conditions tested can be found in Supplementary Table 1. Our experimental design and results followed the convention that a <u>**positive latency**</u> means the periphery was stimulated **after** the cortex by that time difference. These stimulation offsets lead to physiological offsets calculated at the levels of the spinal cord and cortex; positive latencies result in pre-synaptic activity that preceded post-synaptic activity at the specified location.

### PAS Intervention

We used a PAS protocol of 300 paired stimulations to the motor cortex and designated peripheral muscle, using single pulses of biphasic electrical stimulation 0.2 ms in duration, separated by 0.5 Hz. We note that this is on the higher end in terms of number of paired stimulations compared to previous protocols, and is delivered at a higher frequency – but we reasoned, in the absence of evidence otherwise, that any effect that may be present due to paired stimulation should be enhanced using this slightly more intensive protocol.

Cortical stimulation intensity was set at 1.25 times the threshold for a MEP and muscle stimulation at 1.5 times the threshold to elicit a visible twitch. Thresholds were operationally defined as the minimal stimulation intensity required to induce a response more than 50% of the time. All PAS experiments were completed in the animals’ home cage with a modified cover that enabled us to pass the tethering cable, during free behavior (consisting mostly of walking, grooming, and exploring, sometimes sleeping).

### MEP measurement

Raw EMG data was saved and processed offline in LabChart Version 7 and custom scripts written in MATLAB (codebase at https://github.com/ethierlab/PAS). We plotted all individual responses for each cortical stimulation and manually excluded trials for which there was significant excessive movement artifact, and/or lack of EMG signal (this was rare and the most likely reason was due to an intermittent connection with a faulty cable, repaired or replaced promptly). The resulting set of verified MEPs for each probe were collected for further analysis (Figure 4).

**Figure 4:**
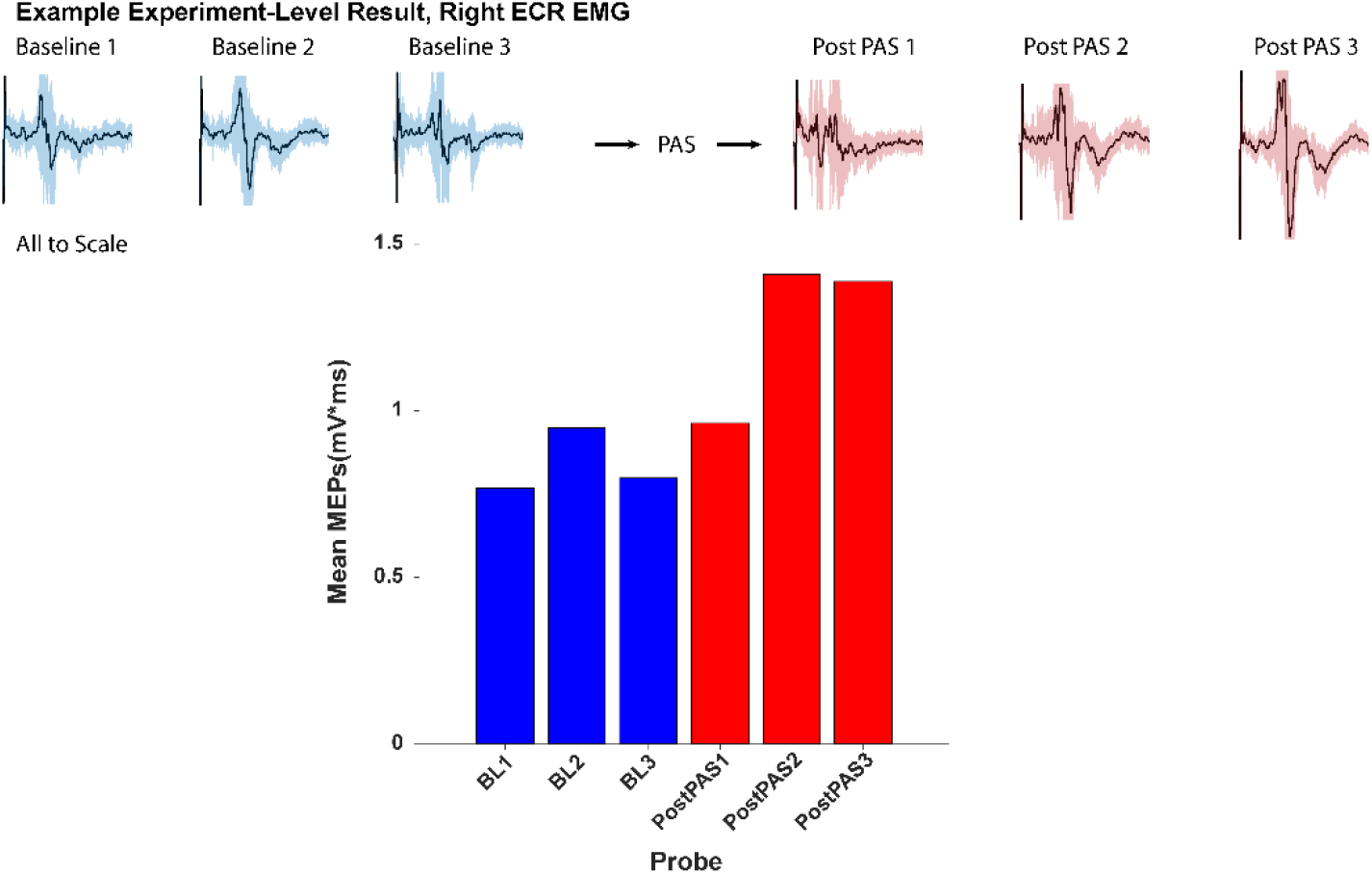
**Representative Experiment-Level Result, MEPs** recorded in the Right ECR of one rat, obtained from electrical cortical stimulation. Shaded area in blue indicates 1 SD about the mean.

MEP amplitudes were initially quantified with three different methods: (1) the peak-to-peak value of individual EMG responses (the literature standard), (2) the mean value of the integral of individual rectified EMG responses, measured over a tailored time window following stimulation, and (3) the integral of the averaged rectified EMG responses, over the same time window. Every individual response to cortical stimulation was first manually screened to exclude any EMG traces containing large movement artifacts or other obvious contamination. In pilot analyses (unpublished data) we assessed qualitatively that the calculation method did not much impact the normalized changes in the MEPs, so we proceeded with taking the integral of the average response for the probe (method 3 above). We reasoned that this approach was most effective in capturing both unimodal and multimodal MEP responses. This decision was made prior to the pooled study data analysis. In summary, the MEP values reported here were thus calculated by first rectifying the filtered EMG signal, then averaging the activity from all stimuli within a probe post-screening, and then calculating the integral of the resulting signal (Figure 4).

### Statistical Analysis

Statistical analysis and visualization were completed using SAS Software, version 9.4 for Windows, and Minitab 18 for Windows. We completed a mixed design ANOVA with repeated measures *on the normalized data* to test the main effects of ISI Timing Condition (CONDITION) with fifteen levels (one for each of 11 timings and four control conditions) as well as PAS Probe (SESSION) with three levels (2, 17 and 32 min after PAS). We also tested for any interaction effects between CONDITION and SESSION. A random effect on the rat was used to account for the randomized block design. The level of significance for the mixed ANOVA was fixed at p < 0.05. Type III Fixed Effects are reported in Table 1, obtained through the Restricted Maximum Likelihood (REML) Estimation method. *Post-hoc* pairwise comparisons between ISI conditions were completed using Dunnett’s Test with the family-wise error rate also set at 0.05. Data from one rat in the ISI +6 condition was removed from the analysis because of poor data quality (very few MEPs in each probe). Normality was assessed on standardized residuals using graphical methods.

## Acknowledgements

We thank Sergiu Ftomov for manufacturing the cortical electrode arrays used for the chronic experiments, and Derek Burns for coding the MATLAB software used for closed-loop recording and stimulation. We are indebted to Chantal Mérette and Geneviève Picher for statistical consultation and discussions throughout the entire study process. We also thank Martin Deschênes and Maxime Demers for sharing laboratory space and equipment with us, and their practical advice on several key experiments, as well as Martin’s and Michaël Elbaz’s feedback on the manuscript. Finally, we are indebted the team in the animal facility at Centre de Recherche CERVO for their continual care of the animals used in the experiments for the duration of this study.

## Author Contributions

WT wrote the first draft of the manuscript. WT, MHL and CE conducted the experiments. WT and CE designed the study, analyzed data and critically revised subsequent drafts to the manuscript. MHL provided critical revisions to the manuscript for intellectual content. All authors approved the final version of the paper for submission.

---

**Supplementary Figure 1:**
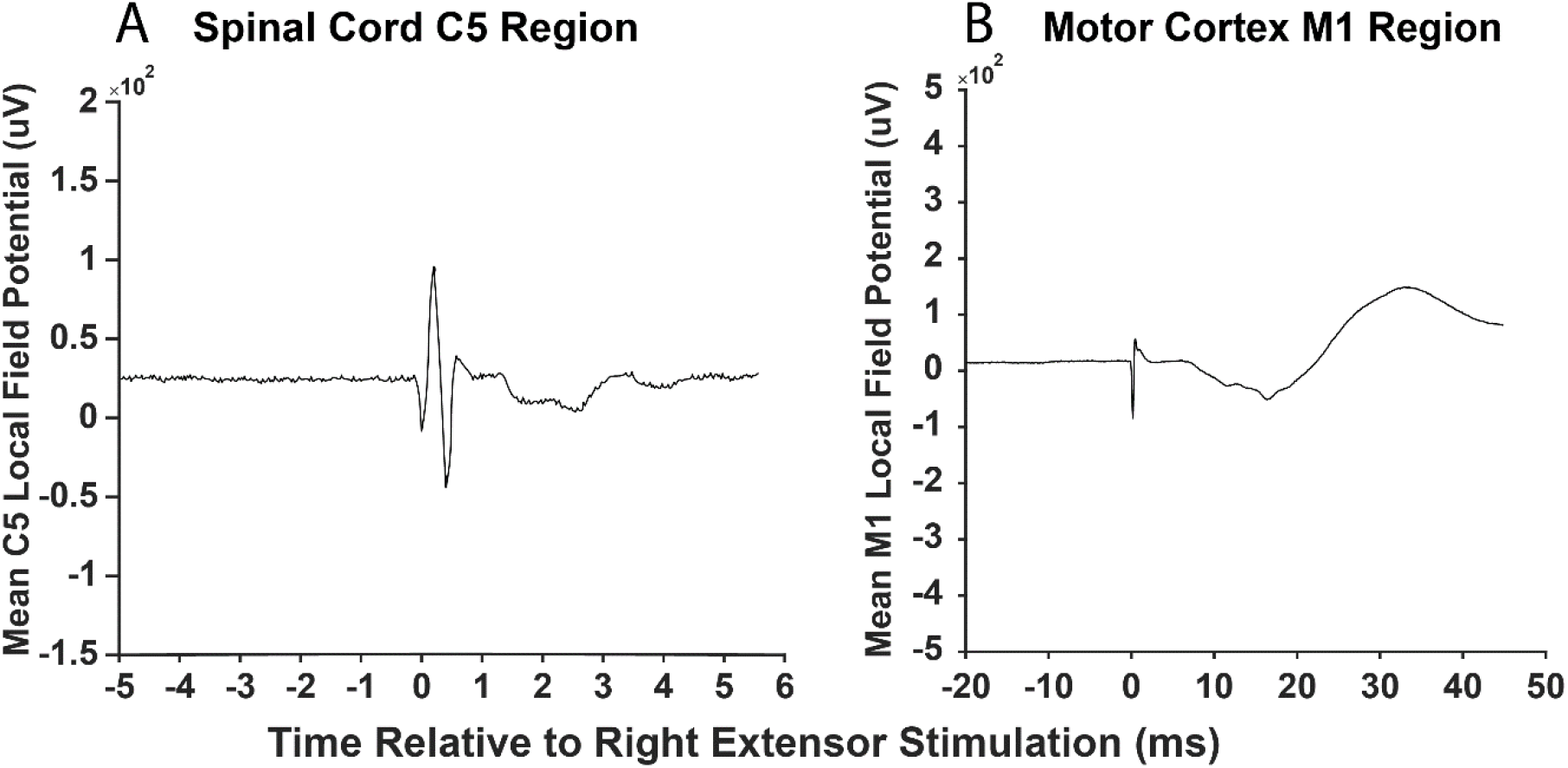
**Spinal and Cortical Evoked Potentials from Peripheral Stimulation,** demonstrating in **A:** the latency of the deepest trough response at 3 ms in the C5 region of the spinal cord, averaged across 200 stimulations, and in **B:** measuring the latency of the deepest trough response at 16 ms in the cortex, averaged across 360 stimulations.

**Supplementary Table 1:** **Study Design for Chronic Experiments** and *a priori* Randomization Order (number within each cell). The final sample sizes for each PAS condition are written to the left. Sessions used in the final dataset are shaded in grey.

| Chronic PAS Experimental Matrix |  |  |  |  |  |  |  |  |  |  |
| --- | --- | --- | --- | --- | --- | --- | --- | --- | --- | --- |
| Condition | Rat |  |  |  |  |  |  |  |  |  |
|  | 1 | 2 | 3 | 4 | 5 | 6 | 7* | 8 | 9 | 10 |
| **ISI Condition** |  |  |  |  |  |  |  |  |  |  |
| STDP -23 (N = 7) | 14 | 14 | 3 | 1 | 3 | 10 | 3 | 11 | 4 | 1 |
| STDP -33 (N = 8) | 9 | 2 | 1 | 11 | 6 | 9 | 4 | 15 | 14 | 15 |
| STDP -48 (N = 7) | 7 | 4 | 6 | 6 | 11 | 7 | 13 | 5 | 9 | 6 |
| STDP -15 (N = 5) | 15 | 15 | 15 | 15 | 15 | 15 | 2 | 3 | 1 | 14 |
| STDP -10 (N = 6) | 12 | 1 | 12 | 14 | 7 | 12 | 8 | 10 | 11 | 10 |
| STDP 0 (N = 6) | 1 | 12 | 11 | 12 | 13 | 14 | 5 | 2 | 10 | 8 |
| STDP +6 (N = 8) | 2 | 6 | 9 | 7 | 9 | 5 | 7 | 9 | 5 | 13 |
| STDP +10 (N = 8) | 6 | 9 | 2 | 4 | 14 | 6 | 15 | 13 | 6 | 5 |
| STDP +12 (N = 7) | 10 | 5 | 13 | 10 | 12 | 4 | 1 | 8 | 7 | 12 |
| STDP +15 (N = 7) | 8 | 11 | 5 | 3 | 1 | 1 | 12 | 14 | 13 | 9 |
| STDP +25 (N = 7) | 13 | 8 | 7 | 13 | 4 | 3 | 14 | 4 | 12 | 11 |
| STDP Timing Control +505 (N = 6) | 11 | 3 | 8 | 9 | 2 | 13 | 9 | 12 | 15 | 7 |
| **STDP Controls** |  |  |  |  |  |  |  |  |  |  |
| Cortex Only Stimulation (Cx) (N = 7) | 5 | 7 | 14 | 5 | 10 | 2 | 10 | 1 | 3 | 2 |
| Muscle Only Stimulation (Ms) (N = 6) | 3 | 10 | 4 | 8 | 5 | 11 | 11 | 7 | 2 | 3 |
| No Stimulation Control (No) (N = 8) | 4 | 13 | 10 | 2 | 8 | 8 | 6 | 6 | 8 | 4 |

\* Implant Failure Prevented Data Collection

**Supplementary Table 2:** **Muscle Distribution** (ISI Conditions, Latencies are Stimulation Offsets). * = PAS muscle stimulation component was between one EMG electrode and reference.

| Rat/Muscle | *Extensor Carpi Radialis* | *Biceps* | *Trapezius* |
| --- | --- | --- | --- |
| Rat 1 | -10, 0, +6, +10,<br>+12, +15, +25,<br>+505.<br>-48, -33, -23.<br>Ms, No, Cx. | --- | --- |
| Rat 2 | --- | --- | -10, 0, +6, +10,<br>+12, +15, +25,<br>+505*.<br>-15, -23, -33*,<br>-48*.<br>Cx, Ms, No. |
| Rat 3 | +6 | --- | +10<br>-33 |
| Rat 4 | -23<br>No |  |  |
| Rat 5 | --- | --- | -10, 0, +6, +10,<br>+12, +15, +25,<br>+505.<br>-15, -23, -33, -48.<br>Cx, Ms, No. |
| Rat 6 | +6*, +10*, +12*,<br>+15*, +25*.<br>-33*, -48*.<br>Cx*, No*. | --- | --- |
| Rat 7 (implant failure before useful data could be collected) | --- | --- | --- |
| Rat 8 | -10*, -0*, +6*,<br>+10*, +12*, +15*,<br>+25*, +505*.<br>-15*, -23*, -33*, | --- | --- |
|  | -48*.<br>Cx, Ms*, No*. |  |  |
| Rat 9<br>(NB: Manual<br>Range Translation<br>During Closed<br>Loop Due to<br>Heartbeat). | --- | -10*, 0*, +6*,<br>+10*, +12*, +15*,<br>+25*, +505*.<br>-15*, -23*, -33*, -<br>48*.<br>Cx*, Ms*, No*. | --- |
| Rat 10 | -10, 0, +6, +10,<br>+12, +15, +25.<br>-15, -23, -33, -48.<br>Cx, Ms, No. | --- | --- |

## Works Cited

Abraira, V. E. and D. D. Ginty (2013). “The sensory neurons of touch.” Neuron 79(4): 618–639.

Alstermark, B., J. Ogawa and T. Isa (2004). “Lack of monosynaptic corticomotoneuronal EPSPs in rats: disynaptic EPSPs mediated via reticulospinal neurons and polysynaptic EPSPs via segmental interneurons.” J Neurophysiol 91(4): 1832–1839.

Bi, G. Q. and M. M. Poo (1998). “Synaptic modifications in cultured hippocampal neurons: dependence on spike timing, synaptic strength, and postsynaptic cell type.” J Neurosci 18(24): 10464–10472.

Bunday, K. L. and M. A. Perez (2012). “Motor recovery after spinal cord injury enhanced by strengthening corticospinal synaptic transmission.” Curr Biol 22(24): 2355–2361.

Bunday, K. L., M. A. Urbin and M. A. Perez (2018). “Potentiating paired corticospinal-motoneuronal plasticity after spinal cord injury.” Brain Stimul 11(5): 1083–1092.

Carson, R. G. and N. C. Kennedy (2013). “Modulation of human corticospinal excitability by paired associative stimulation.” Front Hum Neurosci 7: 823.

Castel-Lacanal, E., A. Gerdelat-Mas, P. Marque, I. Loubinoux and M. Simonetta-Moreau (2007). “Induction of cortical plastic changes in wrist muscles by paired associative stimulation in healthy subjects and post-stroke patients.” Exp Brain Res 180(1): 113–122.

Castel-Lacanal, E., P. Marque, J. Tardy, X. de Boissezon, V. Guiraud, F. Chollet, I. Loubinoux and M. S. Moreau (2009). “Induction of cortical plastic changes in wrist muscles by paired associative stimulation in the recovery phase of stroke patients.” Neurorehabil Neural Repair 23(4): 366–372.

Cho, W., N. Sabathiel, R. Ortner, A. Lechner, D. C. Irimia, B. Z. Allison, G. Edlinger and C. Guger (2016). “Paired Associative Stimulation Using Brain-Computer Interfaces for Stroke Rehabilitation: A Pilot Study.” Eur J Transl Myol 26(3): 6132.

Cooper, S. J. (2005). “Donald Of Hebb’s synapse and learning rule: a history and commentary.” Neurosci Biobehav Rev 28(8): 851–874.

Darling, W. G., S. L. Wolf and A. J. Butler (2006). “Variability of motor potentials evoked by transcranial magnetic stimulation depends on muscle activation.” Exp Brain Res 174(2): 376–385.

Elger, C. E., E. J. Speckmann, H. Caspers and R. W. Janzen (1977). “Cortico-spinal connections in the rat. I. Monosynaptic and polysynaptic responses of cervical motoneurons to epicortical stimulation.” Exp Brain Res 28(3-4): 385–404.

Feldman, D. E. (2012). “The spike-timing dependence of plasticity.” Neuron 75(4): 556–571.

Ferris, J. K., J. L. Neva, B. A. Francisco and L. A. Boyd (2018). “Bilateral Motor Cortex Plasticity in Individuals With Chronic Stroke, Induced by Paired Associative Stimulation.” Neurorehabil Neural Repair 32(8): 671–681.

Grospretre, S., N. Gueugneau, A. Martin and R. Lepers (2017). “Central Contribution to Electrically Induced Fatigue depends on Stimulation Frequency.” Med Sci Sports Exerc 49(8): 1530–1540.

Hebb, D. (1949). The Organization of Behavior, Wiley.

Hennings, K., E. N. Kamavuako and D. Farina (2007). “The recruitment order of electrically activated motor neurons investigated with a novel collision technique.” Clin Neurophysiol 118(2): 283–291.

Hori, N., J. S. Carp, D. O. Carpenter and N. Akaike (2002). “Corticospinal transmission to motoneurons in cervical spinal cord slices from adult rats.” Life Sci 72(4-5): 389–396.

Huang, L. and G. Yang (2015). “Repeated exposure to ketamine-xylazine during early development impairs motor learning-dependent dendritic spine plasticity in adulthood.” Anesthesiology 122(4): 821–831.

Kujirai, K., T. Kujirai, T. Sinkjaer and J. C. Rothwell (2006). “Associative plasticity in human motor cortex during voluntary muscle contraction.” J Neurophysiol 96(3): 1337–1346.

Liang, F. Y., V. Moret, M. Wiesendanger and E. M. Rouiller (1991). “Corticomotoneuronal connections in the rat: evidence from double-labeling of motoneurons and corticospinal axon arborizations.” J Comp Neurol 311(3): 356–366.

Markram, H., J. Lubke, M. Frotscher and B. Sakmann (1997). “Regulation of synaptic efficacy by coincidence of postsynaptic APs and EPSPs.” Science 275(5297): 213–215.

McGie, S. C., K. Masani and M. R. Popovic (2014). “Failure of spinal paired associative stimulation to induce neuroplasticity in the human corticospinal tract.” J Spinal Cord Med 37(5): 565–574.

McKay, W. B., S. M. Tuel, A. M. Sherwood, D. S. Stokic and M. R. Dimitrijevic (1995). “Focal depression of cortical excitability induced by fatiguing muscle contraction: a transcranial magnetic stimulation study.” Exp Brain Res 105(2): 276–282.

Mishra, A. M., A. Pal, D. Gupta and J. B. Carmel (2017). “Paired motor cortex and cervical epidural electrical stimulation timed to converge in the spinal cord promotes lasting increases in motor responses.” J Physiol 595(22): 6953–6968.

Mishra, A. M., A. Pal, D. Gupta and J. B. Carmel (2017). “Paired motor cortex and cervical epidural electrical stimulation timed to converge in the spinal cord promotes lasting increases in motor responses.” J Physiol.

Muller-Dahlhaus, J. F., Y. Orekhov, Y. Liu and U. Ziemann (2008). “Interindividual variability and age-dependency of motor cortical plasticity induced by paired associative stimulation.” Exp Brain Res 187(3): 467–475.

Palmer, J. A., S. L. Wolf and M. R. Borich (2018). “Paired associative stimulation modulates corticomotor excitability in chronic stroke: A preliminary investigation.” Restor Neurol Neurosci 36(2): 183–194.

Pitcher, J. B. and T. S. Miles (2002). “Alterations in corticospinal excitability with imposed vs. voluntary fatigue in human hand muscles.” J Appl Physiol (1985) 92(5): 2131–2138.

Rogers, L. M., D. A. Brown and J. W. Stinear (2011). “The effects of paired associative stimulation on knee extensor motor excitability of individuals post-stroke: a pilot study.” Clin Neurophysiol 122(6): 1211–1218.

Saito, K., K. Sugawara, S. Miyaguchi, T. Matsumoto, H. Kirimoto, H. Tamaki and H. Onishi (2014). “The modulatory effect of electrical stimulation on the excitability of the corticospinal tract varies according to the type of muscle contraction being performed.” Front Hum Neurosci 8: 835.

Sale, M. V., M. C. Ridding and M. A. Nordstrom (2007). “Factors influencing the magnitude and reproducibility of corticomotor excitability changes induced by paired associative stimulation.” Exp Brain Res 181(4): 615–626.

Shin, H. I., T. R. Han and N. J. Paik (2008). “Effect of consecutive application of paired associative stimulation on motor recovery in a rat stroke model: a preliminary study.” Int J Neurosci 118(6): 807–820.

Song, S., K. D. Miller and L. F. Abbott (2000). “Competitive Hebbian learning through spike-timing-dependent synaptic plasticity.” Nat Neurosci 3(9): 919–926.

Stefan, K., E. Kunesch, L. G. Cohen, R. Benecke and J. Classen (2000). “Induction of plasticity in the human motor cortex by paired associative stimulation.” Brain 123 Pt 3: 572–584.

Stefan, K., M. Wycislo and J. Classen (2004). “Modulation of associative human motor cortical plasticity by attention.” J Neurophysiol 92(1): 66–72.

Suppa, A., A. Quartarone, H. Siebner, R. Chen, V. Di Lazzaro, P. Del Giudice, W. Paulus, J. C. Rothwell, U. Ziemann and J. Classen (2017). “The associative brain at work: Evidence from paired associative stimulation studies in humans.” Clin Neurophysiol 128(11): 2140–2164.

Tarri, M., N. Brihmat, D. Gasq, B. Lepage, I. Loubinoux, X. De Boissezon, P. Marque and E. Castel-Lacanal (2018). “Five-day course of paired associative stimulation fails to improve motor function in stroke patients.” Ann Phys Rehabil Med 61(2): 78–84.

Tarri, M., N. Brimhat, D. Gasq, B. Lepage, I. Loubinoux, X. De Boissezon, P. Marque and E. Castel-Lacanal (2018). “Five-day course of paired associative stimulation fails to improve motor function in stroke patients.” Ann Phys Rehabil Med 61(2): 78–84.

Taylor, J. L. and P. G. Martin (2009). “Voluntary motor output is altered by spike-timing-dependent changes in the human corticospinal pathway.” J Neurosci 29(37): 11708–11716.

Tosolini, A. P. and R. Morris (2012). “Spatial characterization of the motor neuron columns supplying the rat forelimb.” Neuroscience 200: 19–30.

Urbin, M. A., R. A. Ozdemir, T. Tazoe and M. A. Perez (2017). “Spike-timing-dependent plasticity in lower-limb motoneurons after human spinal cord injury.” J Neurophysiol 118(4): 2171–2180.

Yagishita, S., A. Hayashi-Takagi, G. C. Ellis-Davies, H. Urakubo, S. Ishii and H. Kasai (2014). “A critical time window for dopamine actions on the structural plasticity of dendritic spines.” Science 345(6204): 1616–1620.

Yang, G., P. C. Chang, A. Bekker, T. J. Blanck and W. B. Gan (2011). “Transient effects of anesthetics on dendritic spines and filopodia in the living mouse cortex.” Anesthesiology 115(4): 718–726.

Zhang, X. Y., Y. F. Sui, T. C. Guo, S. H. Wang, Y. Hu and Y. S. Lu (2018). “Effect of Paired Associative Stimulation on Motor Cortex Excitability in Rats.” Curr Med Sci 38(5): 903–909.

